# Recurrent Hermaphroditism and Sex-Biased ABCDE Gene Expression Reveal Latent Floral Plasticity in the Pedunculate Oak Lineage (*Q. robur* s.l.)

**DOI:** 10.64898/2026.07.14.738412

**Authors:** Hugo R. Afonso, Mariana Macedo, Herlander Azevedo, Carlos Vila-Viçosa, M. Manuela R. Costa

**Affiliations:** CBMA (Centre of Molecular and Environmental Biology), University of Minho, Campus de Gualtar, 4710 -057 Braga, Portugal; BIOPOLIS Program in Genomics, Biodiversity and Land Planning, CIBIO, Campus de Vairão, 4485 -661 Vairão, Portugal; CIBIO, Research Centre in Biodiversity and Genetic Resources, InBIO Associate Laboratory, Campus de Vairão, Universidade do Porto, 4485-661 Vairão, Portugal; Department of Biology, Faculty of Sciences, University of Porto, 4169-007 Porto, Portugal; MHNC-UP—Museu de História Natural e da Ciência da Universidade do Porto – Herbário PO, Universidade do Porto, Praça Gomes Teixeira, 4099-002 Porto, Portugal

**Keywords:** ABCDE Model, Flower organ development, Hermaphroditic flowers, Homeotic genes, Monoecy, *Quercus orocantabrica*

## Abstract

**Background and Aims:** The development of unisexual flowers relies on the tight coordination of flower organ identity and sex determination. The genus *Quercus* is typically considered strictly monoecious, bearing fully segregated male and female flowers within the same individual tree. However, several reports of atypical flowering across the genus challenge this canonical view, suggesting that flowering in oaks may be more flexible than traditionally assumed. In this work, the dynamics of flower development in *Quercus orocantabrica* were examined to correlate contrasting floral morphologies with divergent molecular profiles.

**Methods:** The flowering phenology of *Q. orocantabrica* trees was closely monitored over several individuals and years, together with a detailed floral morphological analysis of male, female and atypical flowers. Key floral homeotic gene homologues were identified, and their expression assayed in the development of different flowers.

**Key Results:** Recurrent and widespread hermaphroditic flowering was detected in several *Q. orocantabrica* trees, frequently associated with unseasonal flowering events. Gene expression analysis of male, female and hermaphroditic flowers revealed a sex-biased expression of *Q. orocantabrica* B- and C-class genes, with the B-class gene *QoPI* in particular being tightly associated with the presence of fully-developed stamens. In addition, the expression of the C-class gene *QoSHP* contrasted with reports in other Fagaceae, highlighting a potential functional divergence of the C/D-class lineage within the family.

**Conclusions:** The results here depicted indicate that the dynamics of floral sex identity in oaks are more plastic than traditionally assumed, supporting a reinterpretation of oak reproductive biology based on a versatile and resilient framework responsive to different developmental contexts.

## 1 Introduction

The development of separate unisexual flowers has emerged several times throughout evolution, relying on the tight regulation of flower organ initiation and identity (Barrett, 2002; Mitchell and Diggle, 2005). Floral unisexuality is found in approximately 10 % of angiosperms, serving as a main driver of cross-breeding and gene flow, and acts as the main reproductive strategy in some plant families, including the predominantly monoecious Fagaceae (Charlesworth and Charlesworth, 1978; Kubitzki, 1993; Barrett, 2002).

Within the Fagaceae family, the genus *Quercus* L. represents one of the most widespread and ecologically dominant groups of woody angiosperms in the Northern Hemisphere. Its evolutionary history is characterised by extensive morphological variation, weak reproductive barriers, and frequent hybridisation, resulting in a reticulate phylogenetic structure often described as an oak syngameon (Burger, 1975; Denk *et al*., 2017; Hipp *et al*., 2020). The reproductive biology of *Quercus* has traditionally been considered highly conserved. Oaks are described as monoecious, with functionally unisexual flowers arranged in separate inflorescences (staminate catkins and pistillate spikes) (Kaul, 1985; Franco, 1990; Nixon, 1993). This condition is regarded as a derived state within the Fagaceae, associated with developmental specialization towards anemophilous pollination (Kaul, 1985, 1986). However, despite its prominence, oak flowering has historically received little attention from botanists, as it is often considered of reduced taxonomic value (Romero *et al*., 2007). As a result, the genus has been widely treated as bearing strictly unisexual flowers, an assumption that has remained largely unchallenged in both systematic and ecological contexts.

Despite this canonical model, reports of hermaphroditic or intersexual flowers have been documented across multiple *Quercus* lineages over the last century. Historic works by Greene (1889) and Camus (1936-1938) already cite the occasional presence of staminodes in female flowers of Californian and South-east Asian oaks. Later, foundational work by Tucker (1972) and collaborators (1980) demonstrated the presence of fully bisexual flowers in North American white oaks, often associated with altered phenological patterns. These flowers contained both viable stamens and functional pistillate structures, indicating true reproductive functionality rather than incomplete or vestigial development (Tucker, 1972; Tucker *et al*., 1980). Subsequent observations confirmed similar patterns in diverse taxa and regions, including Mediterranean species such as *Quercus suber* and *Quercus ilex*, Mexican white oaks and other North American and South-east Asian species (Soepadmo and van Steenis, 1972; Kaul, 1986; Varela and Valdiviesso, 1996; Rangel *et al*., 2000; Borzan and Stabentheiner, 2002; Maqueda, 2007; Romero *et al*., 2007; Borgardt, 2013; Keuter, 2018). In many cases, these occurrences were associated with late-season or secondary flowering, environmental stress, or atypical developmental cycles, suggesting that the sexual behaviour of oaks may be more variable than traditionally assumed (Tucker, 1972; Tucker *et al*., 1980; Borzan and Stabentheiner, 2002; Keuter, 2018). Collectively, these reports indicate that hermaphroditism is not restricted to a particular section or geographic region, but may represent a recurring phenomenon across the genus. Nevertheless, the occurrence of hermaphroditic flowering has not been reported in European pedunculate oaks, particularly within the lineage traditionally associated with *Quercus robur* L., the type species of the genus.

The taxonomic interpretation of the pedunculate oak lineage has been historically complex, particularly within the Northwestern Iberian species circumscription. Here, material that was originally described as *Quercus robur* subsp. *broteroana* O. Schwarz (Schwarz, 1937), has been elevated under the International Code of Nomenclature to the species rank – *Quercus orocantabrica* Rivas Mart. & al. – after a reassessment of Iberian material supported a consistent combination of diagnostic characters (Vila-Viçosa *et al*., 2023). Recent RAD-seq analyses further supported this recognition as a distinct lineage, with Western Iberian pedunculate oaks forming a sister clade to *Q. robur* sensu stricto (Vila-Viçosa *et al*., 2023; Vila-Viçosa *et al*., 2026). Given the central role of pedunculate oaks in defining the reproductive model of the genus, the apparent absence of hermaphroditic flowering is noteworthy; while it may reflect a true rarity of the phenomenon in this lineage, it may also result from historical under-detection, especially in a system where floral morphology has been interpreted within a rigid framework of unisexuality.

The mechanisms of floral development have been extensively studied in model species, culminating in the proposal of the ABCDE floral quartet model (Bowman *et al*., 1989; Coen and Meyerowitz, 1991; Meyerowitz *et al*., 1991; Colombo *et al*., 1995; Angenent and Colombo, 1996; Pelaz *et al*., 2000; Theißen, 2001; Ditta *et al*., 2004; reviewed in Bowman and Moyroud, 2024). According to this model, different combinations of transcription factors coordinate the identity of each floral organ: the expression of A-class (*APETALA1-2*, *AP1-2*) and E-class (*SEPALLATA1-4*, *SEP1-4*) genes specifies sepal identity, while A-, B-(*PISTILLATA*, *PI*; *APETALA3*, *AP3*; and *TOMATO MADS 6*, *TM6*) and E-establishes petals; B-, C- (*AGAMOUS*; *AG*) and E- initiates stamens, and C- and E- originates carpels, with the expression of C, D- (*SHATTERPROOF1-2*; *SHP1-2*) and *SEEDSTICK*; *STK*) and E-class genes specifying ovule identity within the carpel (reviewed in Bowman and Moyroud, 2024). Over the past few decades, an increasing body of evidence suggests that these mechanisms are largely conserved among angiosperms, across different flowering habits, phenologies, and floral morphologies (reviewed in Theißen *et al*., 2016; Bowman and Moyroud, 2024).

In the Fagaceae family, the mechanisms that regulate flowering have been studied in *Q. suber*, *Castanea sativa* and *Castanea mollissima* (Sobral and Costa, 2017; Alhinho *et al*., 2021; Liu *et al*., 2021). In these species, the development of unisexual flowers is dependent on a partial redeployment of the ABCDE model, represented by a sex-biased expression of homeotic genes, non-canonical protein-protein interactions, and suspected sub-functionalization of C/D-class genes (Sobral and Costa, 2017; Alhinho *et al*., 2021; Liu *et al*., 2021). Notably, the expression of the respective *PI* homologue is restricted to *Q. suber* male flowers, while being expressed in *Castanea* female flowers that present underdeveloped staminodes, suggesting its pivotal role in unisexual flower identity. However, despite these advancements, the involvement of floral homeotic genes in other *Quercus* species and its correlation with atypical floral morphologies remain unclear.

In this work, recurrent hermaphroditic flowering is reported in the Iberian pedunculate oak *Q. orocantabrica*. The phenomenon is observed across multiple individuals and years, and was analysed through an integrated approach combining floral morphology, phenology, pollen viability, and gene expression, while aiming to understand the involvement of floral homeotic gene expression in the development of unisexual and hermaphroditic flowers.

## 2 Materials and Methods

### 2.1 Biological Material

For three consecutive years (2023-2025), the phenological cycle of several adult *Q. orocantabrica* trees located in the Minho and Porto regions (Northern Portugal, Iberian Peninsula) was closely monitored, with particular emphasis on the timing of bud burst, male and female inflorescence emergence, subsequent vegetative flushes, and the occurrence of unseasonal flowering events. The floral morphology of male, female and atypical flowers was analysed under a S9 D stereo microscope (Leica) and recorded with an MC170 HD camera (Leica). Flowering branch samples were harvested, press-dried and stored at the Herbarium PO (Museu de História Natural e da Ciência da Universidade do Porto) (specimens PO-V77205 – PO-V72215, PO-V72926 and PO-V72927). Female, male and morphologically hermaphroditic flowers in different stages of development were collected from three selected adult trees located in the Gualtar campus, University of Minho (https://www.icampi.uminho.pt/pt/ambiente/green/; 41.56236104 °N, 8.39401066 °W), immediately frozen in liquid nitrogen, and stored at -80 °C until RNA extraction. Male and hermaphroditic flowers were also collected for pollen germination assays.

### 2.3 Pollen Germination Assays

Pollen from mature anthers of male and morphologically hermaphroditic flowers was collected, transferred to a 15 % (w/v) sucrose solution, and incubated for 24 h in a humid chamber at room temperature. Pollen grains were analysed under a microscope, and considered germinated when the pollen tube surpassed the diameter of the pollen grain. Germination rate was calculated from three technical replicates, with a minimum of 500 pollen grains counted in each replicate.

### 2.4 Histological Analysis

Female and hermaphroditic flowers were collected and immediately fixed in 4 % (w/v) paraformaldehyde in 1x PBS (phosphate buffered saline: 130 mM NaCl, 7 mM Na_2_HPO_4_, and 3 mM NaH_2_PO_4_; pH 7.4) under vacuum infiltration, followed by overnight incubation at 4 °C in the fixative solution. The samples were then rehydrated, cleared and wax embedded according to the protocol by Jackson (1991). Tissue sections with 8 μm thickness were obtained using a microtome (SLEE), mounted in pre-coated poly-L-lysine slide, and stained with toluidine blue. The slides were analysed and photographed under a DM 500 (Leica) microscope.

### 2.5 RNA Extraction

Total RNA was extracted from the selected tissues using the CTAB/LiCl method (Chang *et al*., 1993) with some modifications (Le Provost *et al*., 2007; Serrazina *et al*., 2015). RNA integrity was analysed through agarose gel electrophoresis, and concentration and purity estimated with a ND-1000 Spectrophotometer (Nanodrop). DNAse treatment was performed using DNAse I kit (Grisp), following the manufacturer’s instructions.

### 2.6 RT-qPCR Analysis

For gene expression profiling, cDNA was synthesized from 2 μg of each RNA sample using the NZY First-Strand cDNA Synthesis kit (NZYTech MB40001), following the manufacturer’s instructions. Amplification of cDNA was performed using SsoFast EvaGreen Supermix (BioRad), in 10 μL volume reactions containing 250 nM of each gene-specific primer, and 1 μL of diluted cDNA (1:50). Quantitative real-time PCR (RT-qPCR) was conducted on a CFX96 Touch™ Real-Time PCR Detection System (Bio-Rad, Hercules, CA, USA). RT-qPCR consisted of an initial 3 min denaturing step at 95 °C, followed by 40 cycles composed of a 10 s denaturing step at 95 °C and a 10 s annealing/extension step at 60 °C; a melting curve was obtained at the end of each reaction through a 65 to 95 °C gradient at 5 s intervals. The primers used for each reaction are specified in Table S1. Gene expression analysis was based on three biological and three technical replicates. Due to sample availability restrictions, only two biological replicates were obtained for hermaphroditic flowers. Expression levels were normalized using the Livak calculation method (Livak and Schmittgen, 2001), using *QoPP2AA3* and *QoEF1α* as the reference genes (Marum *et al*., 2012).

### 2.7 Phylogenetic Analysis

BCD-like protein sequences from *Q. robur*, and homologous sequences from other selected species, were obtained by performing a BLASTp search against the NCBI Non-Redundant Protein database (https://www.ncbi.nlm.nih.gov/, accessed on 14 May 2025) using *Q. suber* and *Arabidopsis thaliana* sequences as query. Selected species included: *Quercus lobata*, *C. sativa*, *C. mollissima*, *Fagus crenata*, *Betula pendula*, *Corylus avellana*, *Cucumis sativus*, *Malus domestica*, *Prunus persica*, *Populus trichocarpa*, *Citrus unshiu*, *A. thaliana*, *Vitis vinifera*, *Solanum lycopersicum*, *Antirrhinum majus*, *Petunia hybrida*, *Oryza sativa*, *Amborella trichopoda* and *Pinus radiata*. Sequences were aligned using MUSCLE (Edgar, 2004), manually curated for domain preservation, and distances estimated using the Jones-Taylor-Thornton (JTT) model of evolution. A maximum-likelihood phylogenetic tree was generated using the MEGA 11 software (Tamura *et al*., 2021); statistical support for each node was provided from 1000 bootstrap iterations. The protein accession numbers are shown in Table S2.

## 3 Results

*Quercus* species are generally monoecious, bearing separate male and female flowers within the same individual (Kaul, 1985). While typically considered strictly unisexual, intermediate floral morphologies have been occasionally reported in several species, although this phenomenon is still poorly understood (Tucker, 1972; Tucker *et al*., 1980; Keuter, 2018). Earlier observations by our research group detected atypical flower morphologies in *Q. orocantabrica* trees within the Minho and Porto region of northern Portugal, a trait not previously reported in this lineage. As such, in this work, a detailed phenological and floral morphological analysis of *Q. orocantabrica* was performed. The homologues of key floral homeotic genes were identified, and their expression in flowers with contrasting morphologies was analysed.

### 3.1 Phenological Analysis

For three consecutive years (2023-2025), dormancy break and initial bud burst started in early to mid-February in the majority of the analysed specimens. Prior to bud burst, male inflorescences could already be observed developing within the lateral buds, surrounded by the bud scales (Fig. 1A,B). Immediately upon bud burst, unisexual male catkins with well-defined anthers emerged from the buds (Fig. 1C; Fig. 2A), as well as from the axils of the first few leaves. Female inflorescences emerged from some of the axils of newly formed leaves, 7-9 days after male inflorescence emergence, displaying a mild protandrous habit (Fig. 1D).

**Figure 1:**
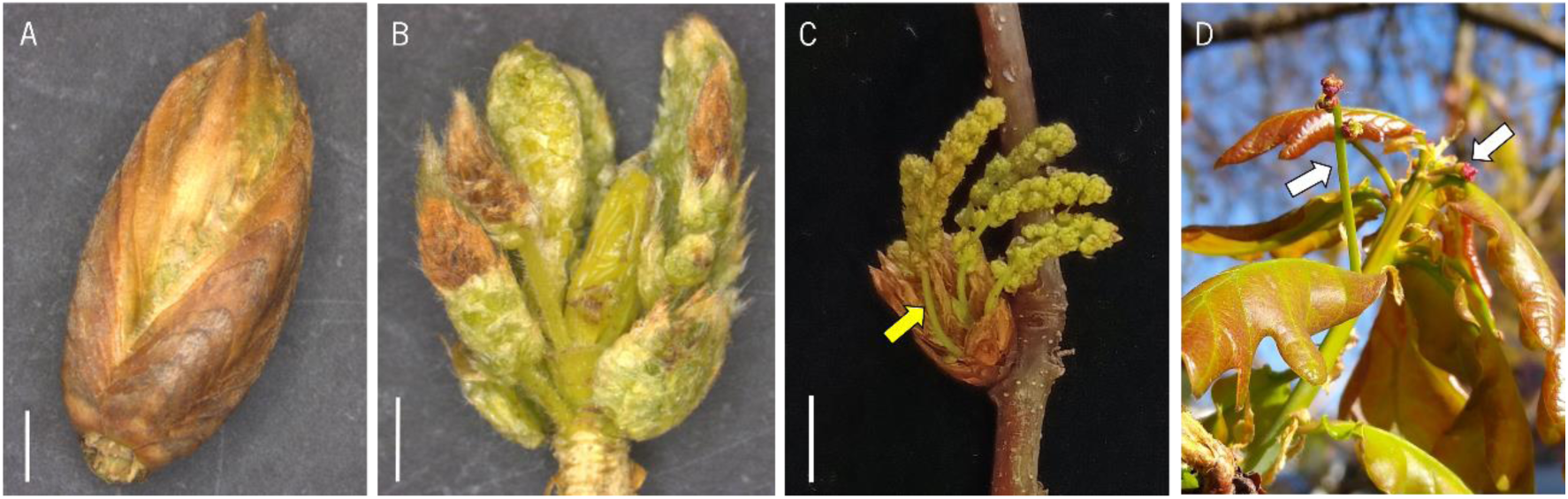
Inflorescence emergence in *Quercus orocantabrica*. **A)** Swollen lateral bud with emerging male inflorescences; **B)** Swollen lateral bud with emerging male inflorescences (scales removed); **C)** Bursted lateral bud with elongating male inflorescences (yellow arrow); **D)** New branch bearing female inflorescences in the axils of new leaves (white arrows). The samples were collected from early February to early March. Scale bars: A, B - 2.5 mm; C - 1 cm.

**Figure 2:**
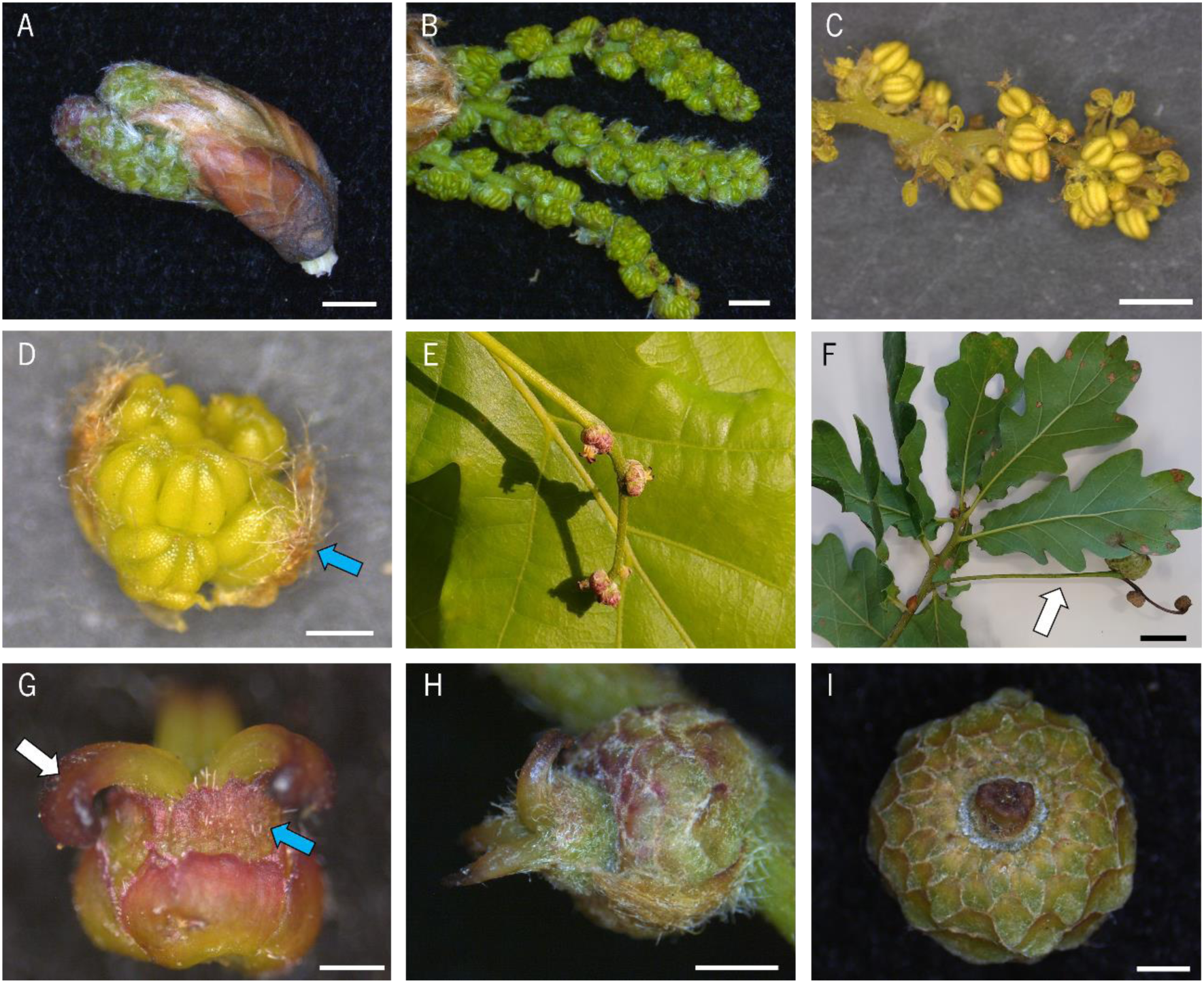
*Quercus orocantabrica* male and female flowers in different stages of development. **A)** Young male catkin emerging from an axillarybud; **B)** Elongating male catkin; **C)** Mature male catkin, flowers shedding pollen; **D)** Mature male flower with visible tepals (blue arrow), prior to pollen shed; **E-F)** Elongated female inflorescence (white arrow); **G)** Young female flower with visible tepals (blue arrow) and receptive stigmas (white arrow); **H-I)** Fertilized female flower in successive stages of development. The samples were collected from early February to late April. Scale bars: D, G : - .5 cm; H, I - 1 cm; A, B, C, F - 2 cm.

To study the development of *Q. orocantabrica* flowers, the morphology of male and female flowers was analysed*. Q. orocantabrica* male flowers were typically arranged in pendant, elongated catkins (Fig. 2B-C), matching what was described for the wider *Quercus* genus (Kaul, 1985, 1986). These flowers typically presented a perianth whorl composed of 4-5 green to brown partially fused tepals, and a stamen whorl bearing 6-8 anthers (Fig. 2D). The female flowers were arranged in a stiff spike with 3-6 flowers (Fig. 2E-F); the female inflorescence presented a long peduncle, characteristic of pedunculate oaks, generally over 8 cm long. Female flowers were composed of a perianth whorl with 5-6 green tepals, often gaining a reddish colour, and a centre whorl bearing 3-4 styles (Fig. 2G). These flowers were partially enclosed by a cupule composed of imbricate concentric scales, which enlarged as the flower matured and the acorn developed (Fig. 2H-I). Overall, these developmental traits followed the established expectations of oak flower biology (Kaul, 1985; Franco, 1990; Nixon, 1993).

Interestingly, several flowers displayed organs with anomalous identities. In some male flowers an additional well-developed anther could be observed on a distinct centre whorl, smaller than the surrounding anthers (Fig. 3A). In addition, some male flowers appeared to display a vestigial, seemingly dysfunctional pistil on the centre whorl (Fig. 3B-C). On some occasions, female inflorescences were found bearing 2-3 apparently normal male flowers instead of female flowers (Fig. 3D-E). Some female flowers also displayed anomalous morphology, presenting stamens with fully developed anthers in a distinct whorl between the styles and the tepals, thereby forming morphologically hermaphroditic flowers (henceforth referred to as Hermaphroditic flowers) (Fig. 3F-I). Most of these flowers appeared to be fertilized, and fruit development progressed normally (Fig. 3J-K). Female flowers bearing aborted stamens were not observed. Hermaphroditic flowers and apparently normal unisexual female flowers were frequently found in the same inflorescence. Some female flowers also appeared to display an atypically high number of styles (up to 10 each); the additional styles were also located on a clearly distinct whorl (Fig. 3L-M). On rare occasions, stamens and styles were found simultaneously on this additional whorl (Fig. 3M). Hermaphroditic flowers with well-developed anthers were found in the majority of the analysed trees throughout the region, indicating that this phenomenon is widespread rather than an isolated event. These flowers were observed over three consecutive years (2023-2025), with most individual trees consistently displaying this trait throughout the study period. An analysed herbarium specimen from the Herbarium PO collection (PO-V 29700!), harvested in Porto in May 1970, was also found to present hermaphroditic flowers (Fig. 3N). This historical specimen extends the temporal record of hermaphroditic flowering in *Q. orocantabrica* by more than five decades, demonstrating that the phenomenon predates the present study and has persisted beyond the recent monitoring period.

**Figure 3:**
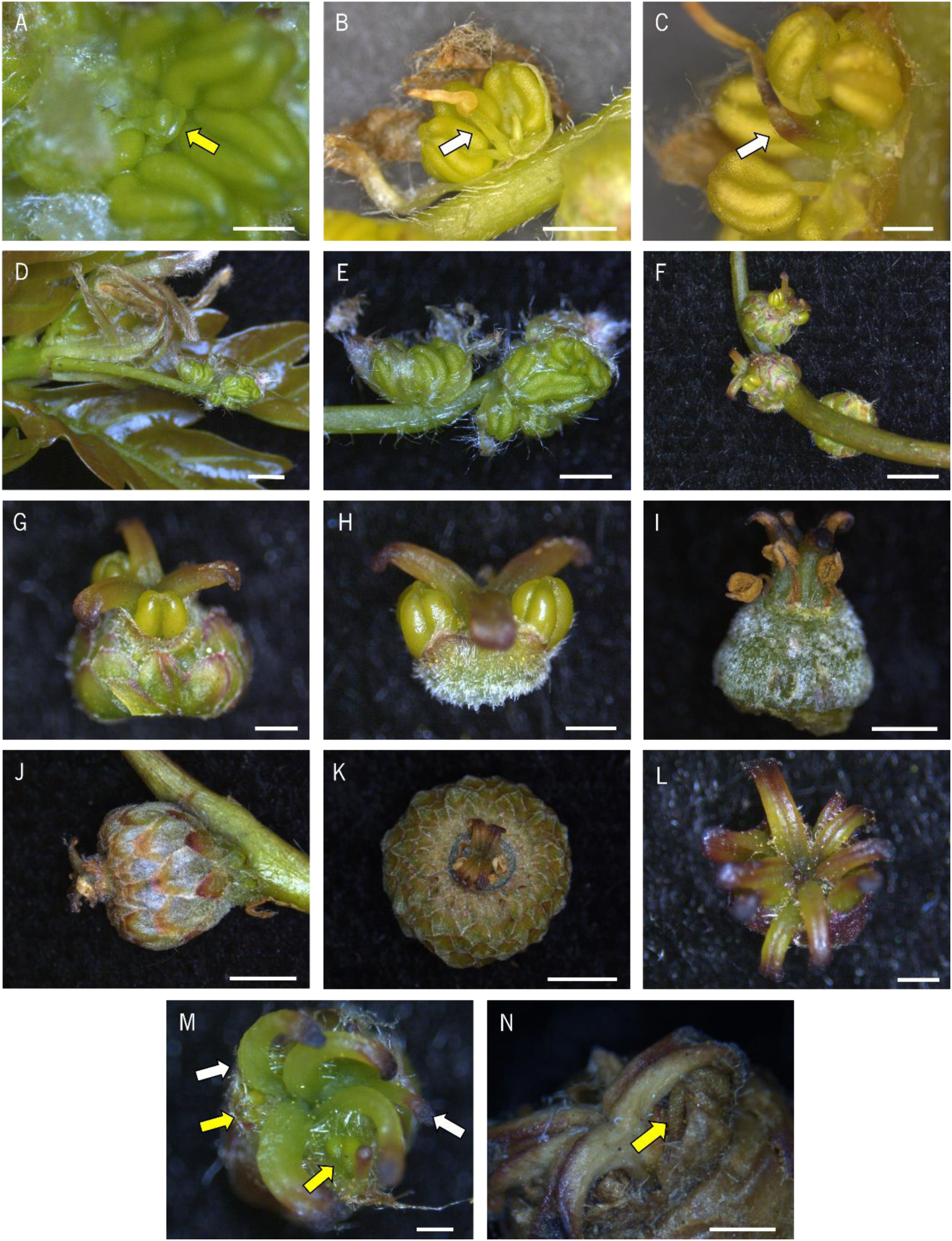
Floral anomalies in *Quercus orocantabrica.* **A)** Male flower bearing an additional stamen on a distinct centre whorl (yellow arrow); **B-C)** Male flowers bearing a non-functional pistil on the centre whorl (white arrow); **D-E)** Female inflorescence bearing male flowers; **F)** Female inflorescence bearing hermaphroditic flowers; **G)** Hermaphroditic flower; **H)** Hermaphroditic flower (cupule removed); **I)** Hermaphroditic flower with stamens in a clearly distinct whorl (cupule and tepals removed); **J-K)** Hermaphroditic flowers progressing through fruit development; **L)** Female flower bearing an abnormal number of styles; **M)** Female flower bearing both styles (white arrows) and stamens (yellow arrows) on a distinct whorl; **N)** Hermaphroditic flower preserved in a herbarium specimen, with a visible stamen (yellow arrow). The samples were collected from early February to mid-May. Scale bars: A, C, G, H, L, N - 0.5 cm; B, E, I, M - 1 cm; D, F, J, K - 2 cm.

*Quercus* species typically display multiple consecutive vegetative flushes over the course of one growth season, in which buds that were formed during the current season experience burst and generate new growth (Natividade, 1950; Kaul, 1986; Bobinac *et al*., 2012). In the analysed *Q. orocantabrica* trees, secondary bud burst was generally observed beginning in mid-to-late May. This new flush did not occur simultaneously within each tree, and it was observed throughout the remainder of the growing season, extending up to late October. In addition, some buds from the previous year that did not burst at the beginning of the season also experienced burst during this time. In 2023 and 2024, the secondary bud burst was followed by unseasonal flowering in a large portion of the analysed trees, with the emergence of new flowers also occurring throughout the remainder of the growth season.

The secondary flowering was composed of three main inflorescence types. From late-May to early-June, apparently normal female inflorescences emerged from the axils of the first few new leaves, bearing both female and hermaphroditic flowers (Fig. 4A). In addition, on rare occasions, female inflorescences were found bearing vegetative structures (buds and leaves) instead of flowers (Fig. 4B-C). From late-May to mid-September, abnormal inflorescences emerged from the bursting buds and the axils of the first few new leaves (Fig. 4D-E), the typical placement of male inflorescences. Both female flowers, hermaphroditic flowers, and apparent male flowers with dysfunctional pistils were found in some of these atypical inflorescences (Fig. 4F-H, respectively). These inflorescences displayed a longer and thicker peduncle, surpassing 10 cm in length, while bearing more abundant flowers than female inflorescences (up to 13 flowers each) (Fig. 4D-E, I). In some cases, the inflorescences appeared to exhibit an acropetal gradient of male identity, with flowers closer to the apex bearing more well-developed anthers (Fig. 4J). Several flowers appeared to be fertilized and to initiate fruit development. However, none reached maturity, as all developing fruits were shed prior to bud set in late autumn. From mid-September to late October, typical male inflorescences bearing normal male flowers emerged from newly-bursted buds; most of these bursted buds did not develop a new branch, and entered senescence shortly after male flower emergence.

**Figure 4:**
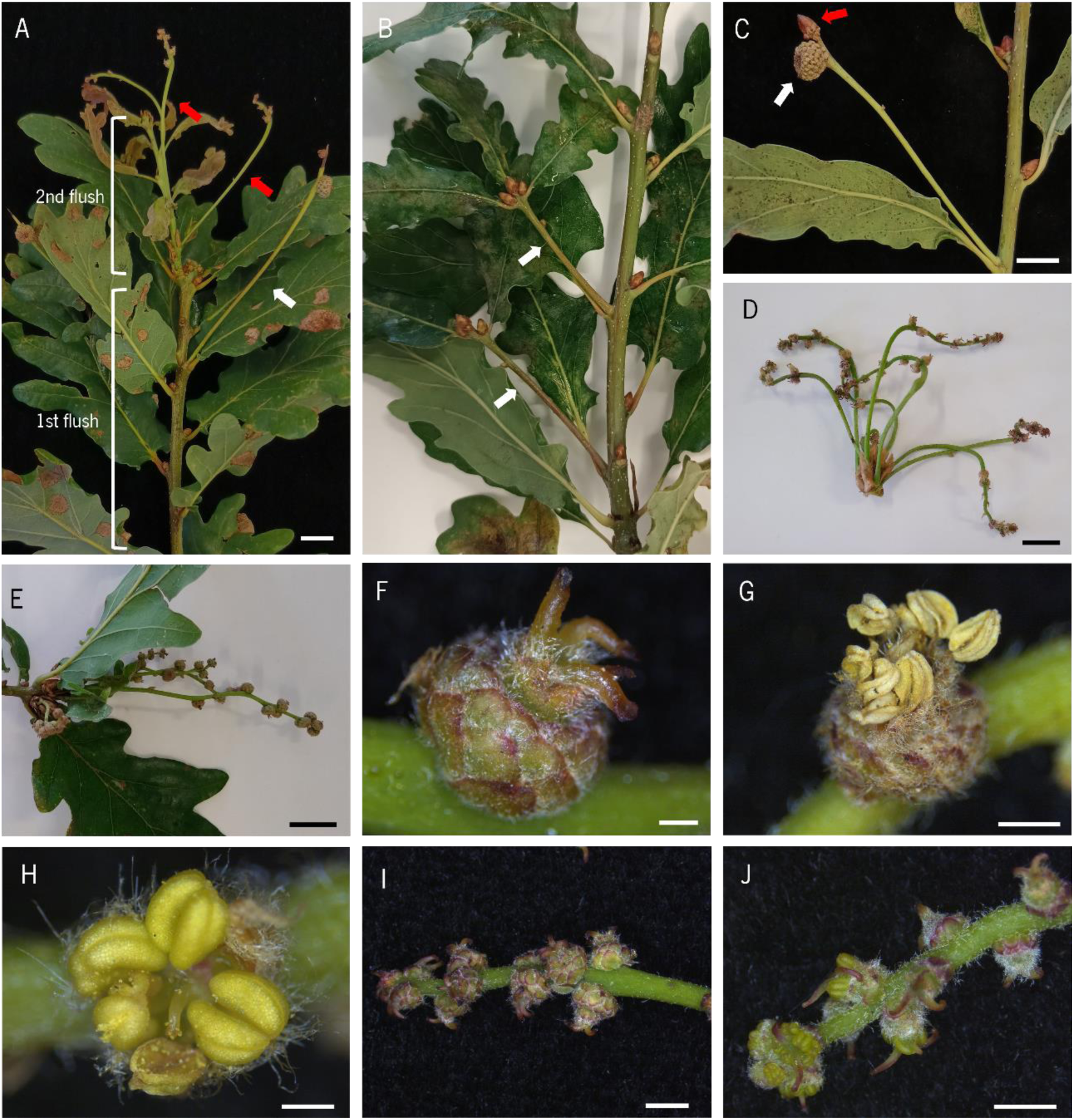
Floral anomalies in the second flowering event of *Quercus orocantabrica.* **A)** Branch displaying two vegetative flushes, each bearing female inflorescences in the axils of new leaves (white arrow: female inflorescence from the first flowering event; red arrows: female inflorescences from the second flowering event); **B)** Female inflorescences bearing vegetative structures; **C)** Female inflorescence bearing both a female flower (white arrow) and a lateral bud (red arrow); **D-E)** Abnormal inflorescences emerging from the bud; **F)** Female flower; **G)** Hermaphroditic flower; **H)** Male flower bearing a dysfunctional pistil; **I)** Abnormal inflorescence bearing 12 female flowers; **J)** Hermaphroditic flowers close to the apical end, bearing more prominent stamens. The samples were collected from late May to late August. Scale bars: F, H - 0.5 cm; C, D, G - 1 cm; A, E, I, J - 2 cm.

To better understand the anatomy of female and hermaphroditic flowers, a histological analysis was conducted. In female flowers, no underdeveloped or aborted stamen primordia was found underneath the perianth (Fig. 5A). In hermaphroditic flowers, well developed anthers with visible pollen grains were observed (Fig. 5B).

**Figure 5:**
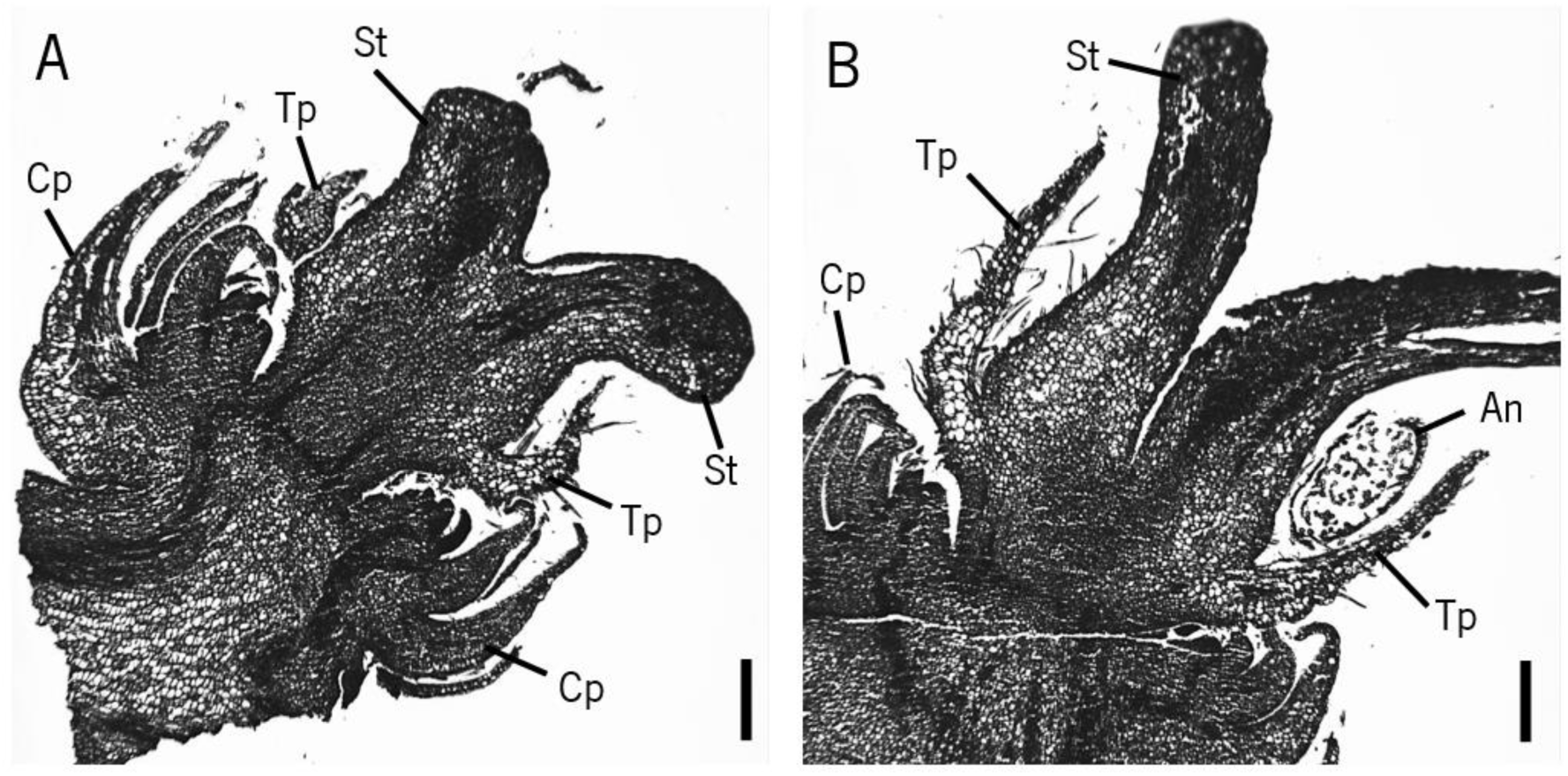
Histological sections of *Quercus orocantabrica* flowers. **A)** Female flower; **B)** Hermaphroditic flower. St, Stigma; Tp, Tepal; An, Anther; Cp, Cupule Scales. Scale bars: 250 μm.

In addition, to check if the anthers from hermaphroditic flowers were functional, pollen from these flowers from both the first and second flowering events was tested for germination capability and compared with pollen from male flowers. Pollen from hermaphroditic flowers from both the first and second flowering events could germinate (Fig. S1), with a rate of 16,46% ± 3,18% and 11,08% ± 9,04%, respectively, which was comparatively lower than pollen derived from male flowers (45,33% ± 2,70%).

### 3.2 Phylogeny of *Quercus* BCD-like genes

Recent studies have explored the role of ABCDE MADS-domain protein homologues in *Q. suber*, *C. sativa* and *C. mollissima* flower development (Sobral and Costa, 2017; Alhinho et al., 2021; Liu *et al*., 2021); however, the conservation of these proteins and their function in other *Quercus* species remains undescribed. While there is currently no genome assembly available for *Q. orocantabrica*, the *Q. robur* genome has been sequenced (Plomion *et al*., 2016). Considering the close genetic proximity of *Q. orocantabrica* and *Q. robur* (Vila-Viçosa *et al*., 2023; Vila-Viçosa *et al*., 2026), the *Q. robur* genome was mined for B-, C- and D-class homologues of the ABCDE model genes. The resulting sequences were used in phylogenetic analysis of the different gene classes. For this purpose, a BLASTp search using *Q. suber* and *A. thaliana* MADS-domain proteins as query was conducted. The queries included PI, AP3 and TM6 (B-class proteins), and AG, SHP1-2 and STK (C/D-class proteins). To investigate the phylogeny of *Q. robur* BCD-like proteins, homologous sequences from species representative of major angiosperm groups were used, together with a basal angiosperm (*Amborella trichopoda*) to root the angiosperm phylogeny and provide evolutionary context prior to the monocot–eudicot split, and a gymnosperm (*Pinus radiata*) as an outgroup. Protein sequences were also compared to analyse the presence and preservation of domains (MADS-, I-, K- and C-terminal) and C-terminal motifs (PI, paleoAP3, euAP3, AG I and AG II).

In each protein query, a single *Q. robur* homologous protein was identified. Protein alignment suggests that the four canonical MIKC-domains are present and conserved in all sequences (Fig. S2). Throughout evolution, the B-class family underwent several duplication events, culminating in three separate lineages: PI, paleoAP3 and euAP3 (Kramer and Irish, 2000). The *Q. robur* orthologues QrPI, QrTM6 and QrAP3 all clustered with the remaining Fagaceae homologues in the respective lineages (Fig. 7A). The canonical PI motif (Kramer *et al*., 1998) is partially conserved in QrPI, and nearly completely conserved within the family (Fig. S3A), with a single amino acid difference in one of the analysed species (*Q. lobata*). The major difference between the paleoAP3 and euAP3 lineages resides in a divergent motif in the C-terminal domain (Kramer *et al*., 1998). The paleoAP3 motif in the TM6 lineage is completely conserved within *Quercus* and *Castanea*, with a single amino acid difference between the two genera (Fig. S3B). Similarly, the euAP3 motif of the analysed Fagaceae sequences displays a partial sequence divergence from the Arabidopsis euAP3; the motif is totally conserved at the family level, with the exception of *Q. lobata*, which displays a complete deletion of the motif (Fig. S3C).

**Figure 7:**
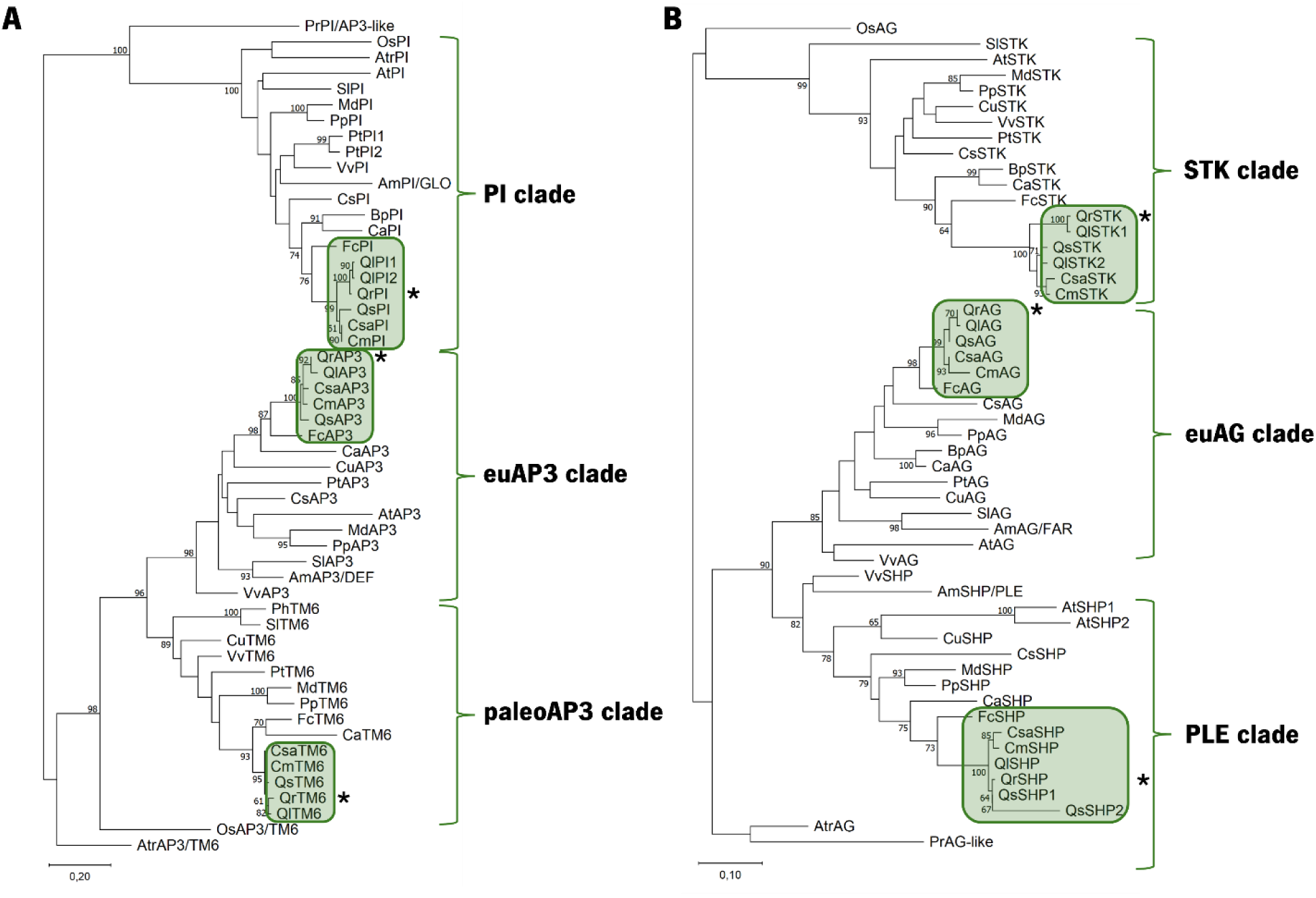
Phylogenetic profiling of the *Quercus robur* BCD protein homologues. **A)** B-class family; **B)** C/D-class family. Evolutionary relationships were determined using the maximum-likelihood method, with 1000 bootstrap replicates. The percentage of replicates in which the taxa clustered together is shown in each node. Fagaceae homologues are highlighted (green boxes). Protein sequences were obtained for the following species: *Quercus robur* (Qr), *Quercus suber* (Qs), *Quercus lobata* (Ql), *Castanea sativa* (Csa), *Castanea mollissima* (Cm), *Fagus crenata* (Fc), *Betula pendula* (Bp), *Corylus avellana* (Ca), *Cucumis sativus* (Cs), *Malus domestic*a (Md), *Prunus persica* (Pp), *Populus trichocarpa* (Pt), *Citrus unshiu* (Cu), *Arabidopsis thaliana* (At), *Vitis vinifera* (Vv), *Solanum lycopersicum* (Sl), *Antirrhinum majus* (Am), *Petunia hybrida* (Ph), *Oryza sativa* (Os), *Amborella trichopoda* (Atr) and *Pinus radiata* (Pr).

The C/D-class subfamily has also experienced multiple duplication events, which led to the divergence of the C and D lineages, and the separation of the former into the euAG and PLE clades (Kramer *et al*., 2004). Both QrAG and QrSHP (C lineage) and QrSTK (D lineage) clustered in the same clade as the other Fagaceae homologues (Fig. 7B). The C/D subfamily is characterised by the presence of the AG motif I and II in the C-terminal domain, which bear some resemblance to the PI and paleoAP3 motifs (Kramer *et al*., 2004). In QrAG, while both motifs are partially conserved, the AG motif II bears a partial deletion (Fig. S3D); these motifs are completely conserved within the Fagaceae. In the PLE lineage, QrSHP has a partially conserved AG motif I, but displays a near-complete deletion of the AG motif II (Fig. S3E). In this protein, the AG motif I is totally conserved within the analysed Fagaceae sequences. By contrast, the *Castanea* genus displays a complete AG motif II in this protein, contrary to the sequences from the *Quercus* genus.

Collectively, these results reveal a high degree of sequence conservation of BCD-like proteins within the Fagaceae family. Considering the close genetic proximity of *Q. robur* and *Q. orocantabrica* and the low degree of sequence divergence in the *Quercus* genus, it is likely that the ABCDE model gene homologues share a similar degree of conservation. Under this presumption, a gene expression analysis during *Q. orocantabrica* flower development was conducted using primer pairs based on the available *Q. robur* gene sequences.

### 3.3 Expression of BCD-like floral homeotic genes in *Q. orocantabrica* flowers

Floral organ development is firmly dependent on a tight temporal and spatial regulation of the expression of MADS-box genes. In *Q. suber* and *C. sativa*, the expression of B and C-class genes was found to be sex-biased, with B-class genes being predominantly expressed in male flowers (Sobral and Costa, 2017; Alhinho *et al*., 2021). To study the involvement of these genes in *Q. orocantabrica* flower development, a quantitative RT-PCR analysis (RT-qPCR) was conducted using cDNA samples from male, female and hermaphroditic flowers from the first flowering event, at different developmental stages. Male flower samples were collected from young inflorescences still concealed within the lateral buds (M0, Fig. 1B), shortly following emergence (M1, Fig. 2A) and during maturation prior to pollen shed (M2, Fig. 2B). Female flower samples were collected after emergence (F1, Fig. 2G), and through the early stages of pollinated flower development (F2 and F3, Fig. 2H-I). Hermaphroditic flower samples were collected in stages equivalent to F1, displaying mature stamens prior to pollen shed, and F2, displaying senescent stamens (H1, Fig. 3G, and H2, Fig. 3J, respectively).

The B-class genes *QoTM6*, *QoAP3*, and *QoPI* displayed higher expression levels in male flowers than in female and hermaphroditic flowers (Fig. 8A-C). While lower than in male flowers, *QoTM6* and *QoAP3* displayed detectable expression levels in female and hermaphroditic flowers. By contrast, *QoPI* expression was not detected in female flowers.

**Figure 8:**
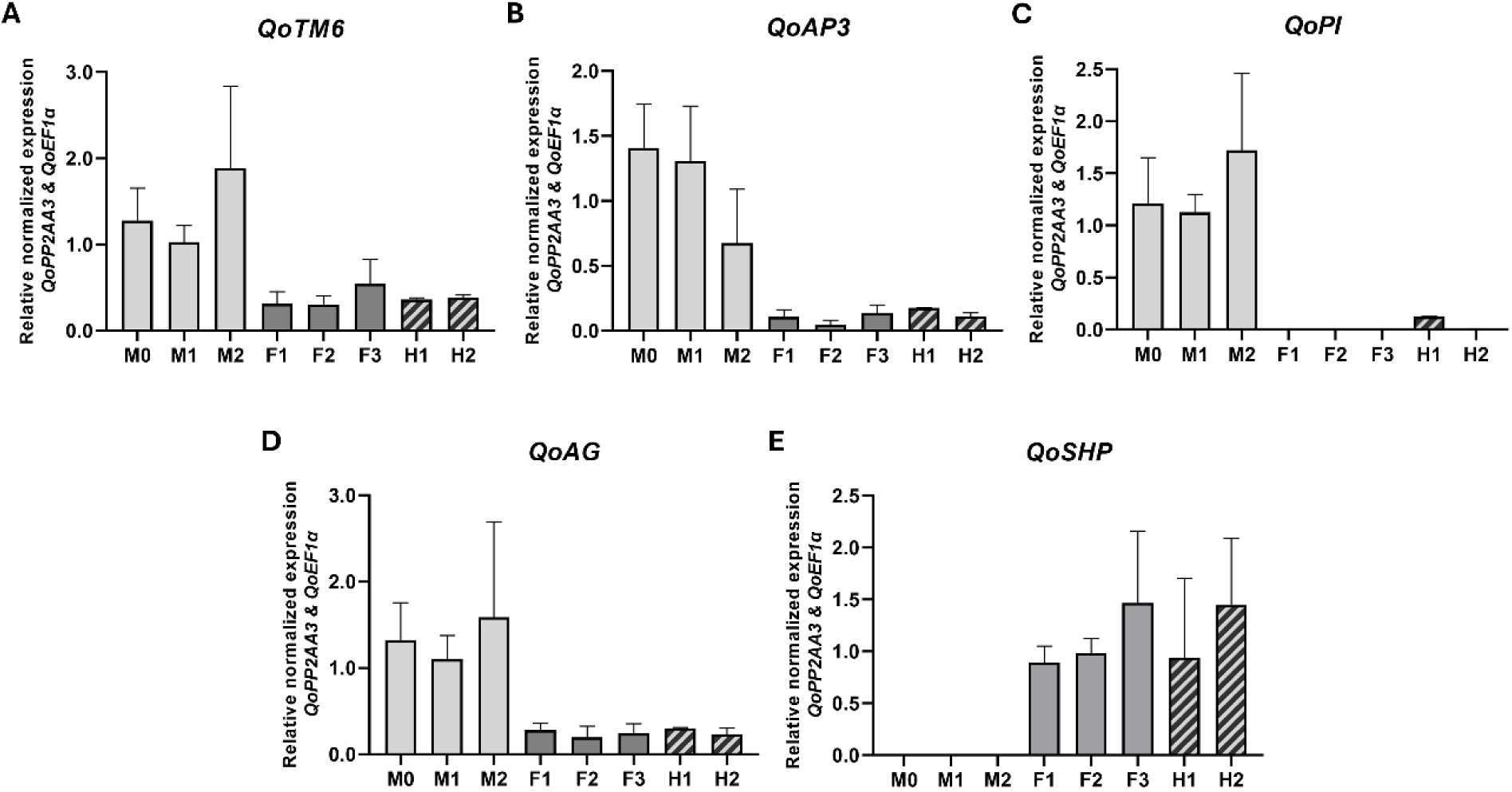
Expression analysis of BCD-like homologous genes in *Quercus orocantabrica* flowers. **A)** *QoTM6*; **B)** *QoAP3*; **C)** *QoPI*; **D)** *QoAG*; **E)** *QoSHP*. Error bars indicate standard deviation of three technical and biological replicates. *QoPP2AA3* and *QoEF1α* were used as reference genes for normalization. M0 – Developing male inflorescence, concealed within the lateral bud; M1 – Young male inflorescence shortly after emergence; M2 – Maturing male inflorescence, male flowers with well-developed stamens prior to pollen shed; F1 – Young female flower; F2-F3 – Fertilized female flower in successive stages of development; H1 – Young hermaphroditic flower with well-developed anthers; H2 – Fertilized hermaphroditic flower, anthers dry following pollen shed.

In hermaphroditic flowers, *QoPI* was only detected at low levels at the H1 stage, prior to anther senescence, suggesting that anther development may be dependent on the expression of this gene.

The C-class gene *QoAG* was found to be more expressed in male flowers than in female and hermaphroditic flowers, in similarity to the analysed B-class genes (Fig. 8D). By contrast, *QoSHP* was not detected in male flowers, being only expressed in female and hermaphroditic flowers (Fig. 8E). The apparent restriction of *QoSHP* expression to female and hermaphroditic flowers suggests that this gene may not be active in the stamen whorl, while possibly maintaining a C/D-function in the centre whorl. Overall, these results show that the expression of BCD-like genes in *Q. orocantabrica* flowers is sex-biased, suggesting a meaningful role in the coordination of unisexual flower identity.

## 4 Discussion

The development of separate unisexual flowers is a trait that promotes crossbreeding and enhances species adaptability and fitness (Charlesworth and Charlesworth, 1978; Barrett, 2002). In the *Quercus* genus, floral development has been traditionally regarded as strictly unisexual, falling upon a fixed phenological sequence (Franco, 1990; Nixon, 1993; Romero, 2007). However, hermaphroditic and unseasonal flowering have been occasionally documented over the past few decades, suggesting that sexual behaviour in oaks may be more flexible than traditionally assumed. In this work, we report frequent and recurrent hermaphroditic flowering in the Iberian pedunculate oak *Q. orocantabrica,* consistently associated with secondary flowering events, a factor that further challenges the canonical interpretation of oak reproductive biology.

The recurrent presence of hermaphroditic flowers across multiple individuals, populations, and years indicates that this condition is not sporadic. This interpretation is further supported by the detection of hermaphroditic flowers in herbarium material collected in 1970, showing that the phenomenon is not limited to the 2023–2025 flowering seasons, and is unlikely to represent a transient abnormality or an artefact of recent environmental conditions. In the present study, hermaphroditic flowers occurred in both primary and secondary flowering phases and displayed a continuum of morphologies ranging from unisexual flowers with vestigial organs to fully bisexual flowers. Similar observations across the genus reinforce the view that hermaphroditism is a recurring feature of oaks, more widespread than the relative shortage of reports may suggest (Tucker, 1972, Tucker *et al*., 1980; Borzan, 2000; Romero *et al*., 2007; Keuter, 2018). Early works proposed that hermaphroditism in *Quercus* may be elicited by atypical environmental conditions, as more recently demonstrated in monoecious mango (*Mangifera indica*) and squash (*Cucurbita pepo*) (Tucker, 1972; Martínez *et al*., 2014; Rajatiya *et al*., 2018). In this framework, hermaphroditism appears not as a rare deviation from a strictly unisexual system, but as a latent and episodic state that occurs under specific developmental and ecological conditions. As such, the reproductive system of oaks is better understood as flexible, rather than a fixed state.

### 4.1 Polycyclism may act as a driver of floral plasticity

*Quercus* species typically display multiple consecutive vegetative flushes over the course of the same growth season, alternating between periods of bud burst and active growth, and small periods of quiescence (Natividade, 1950; Kaul, 1986; Bobinac *et al*., 2012). In temperate *Quercus*, flowering generally only occurs upon the initial spring flush, although occasional unseasonal flowering in succeeding flushes has been reported in several species (Tucker, 1972; Tucker *et al*., 1980; Bobinac *et al*., 2000; Borzan, 2000; Keuter, 2018; García-Mozo *et al*., 2022; Coombes, 2025). In the present study, summer and autumn flowering was detected in several *Q. orocantabrica* trees in different locations of the Minho region. This result contrasts with previous studies with *Q. robur*, in which summer flowering has been reported as a rare occurrence detected in very few individuals (Bobinac *et al*., 2012; Bobinac *et al*., 2000). From an ecological perspective, this phenomenon may be particularly relevant in Iberian environments, where pedunculate oaks frequently experience heterogeneous conditions, including episodic drought, extended growing seasons, and irregular phenological cycles (Amigo *et al*., 2017; Vila-Viçosa *et al*., 2020). In the sympatric *Q. suber*, which can often display both annual and biennial acorn maturation cycles, autumn emergence of female inflorescences reportedly yields viable seed in the following season, reaching maturity in early autumn prior to current-year acorns (Natividade, 1950; Elena-Rossello *et al*., 1993). Thus, the capacity for repeated flushing and secondary flowering may provide a flexible reproductive framework in such systems, allowing these species to take advantage of favourable conditions to increase reproductive output. The associated emergence of hermaphroditic flowers could, in principle, contribute to reproductive assurance when synchrony between male and female phases is disrupted, although the absence of mature fruits from summer and autumn flowers in the present study leaves this potential function unconfirmed. Nevertheless, as no unseasonal flowers were found to reach fruit maturity, the occurrence of summer or autumn flowering in this species might constitute a remnant ancestral trait that fails to provide viable progeny in the current conditions.

The secondary flowering events observed in this study were closely associated with repeated vegetative flushes, forming a continuous developmental sequence linking polycyclic growth to reproductive variation. In other temperate oak species, male and female floral primordia are initiated at different timepoints, a trait thought to underlie the temporal separation of the emergence of both sexes that is characteristically observed in the genus (Merkle *et al*., 1980; Ducousso *et al*., 1993; Sobral *et al*., 2020). Moreover, in *Q. suber*, the separate initiation of floral meristems has been correlated with independent flowering induction events at the molecular level (Sobral *et al*., 2020), which further suggests that typical oak flowering commonly relies in the repeated and successive acquisition of reproductive capability throughout the year.

A significant portion of the inflorescences from secondary flowering events were characterised by irregular architecture, increased floral density, and altered sexual identity, including the co-occurrence of male, female, and hermaphroditic flowers within the same structures. In several cases, an acropetal gradient of sexual identity was observed, suggesting progressive shifts in floral organ specification along the inflorescence axis. Comparable patterns have been reported in other oak species, where late-season flowering results in the emergence of atypical inflorescence architectures and hermaphroditic flowers (Tucker, 1972; Tucker *et al*., 1980; Borzan, 2000; Keuter, 2018; Coombes, 2025). In the Fagaceae family, the coexistence of different flowers within the same inflorescence has been suggested as a primitive intermediate trait in the transition from entomophilous hermaphrodeity to anemophilous monoecy (Kaul, 1986; Ramos, 2007). Taken together, these findings suggest that hermaphroditism and secondary flowering in oaks may reflect an ancestral developmental potential that is rarely expressed under typical conditions but becomes detectable under specific ecological and developmental scenarios.

### 4.2 Unisexual flower development relies on distinct molecular patterns

The development of each floral organ is based on a tight regulation of the floral homeotic genes. In angiosperms, floral organ identity is specified according to the ABCDE model, with different combinations of transcription factors leading to the development of each structure (reviewed in Bowman and Moyroud, 2024). Species that develop separate male and female flowers serve as a good model to study the mechanisms that regulate flower organ development. In this framework, the natural occurrence of floral anomalies provides an additional opportunity to perform studies on gene function, particularly relevant in species where reverse-genetics methods are often unviable. In this work, a gene expression analysis of BCD-like floral homeotic genes in *Q. orocantabrica* male, female and hermaphroditic flowers was performed, revealing a sex-biased expression pattern tightly correlated with floral morphology.

The B-class genes *QoTM6* and *QoAP3* were found to be substantially more expressed in male flowers than female flowers, consistent with previous observations in other Fagaceae (Sobral and Costa, 2017; Alhinho *et al*., 2021; Liu *et al*., 2021). By contrast, *QoPI* expression was not detected in female flowers, being apparently restricted to flowers with developing or well-developed stamens (i.e. male flowers and hermaphroditic flowers prior to anther senescence). In *Q. suber*, *QsPI* was also not found to be expressed in female flowers (Sobral and Costa, 2017). Conversely, the *C. sativa PI* homologue was detected in low levels in female flowers, a trait seemingly correlated with the presence of underdeveloped staminodes underneath the perianth (Alhinho *et al*., 2021). Overall, these data indicate a tight correlation of *PI* expression with the formation of male structures in the Fagaceae, a trend that appears to be conserved in *Q. orocantabrica*. In *A. thaliana*, *pi* knockout mutants fail to initiate stamen organ primordia (Bowman *et al*., 1989, 1991). Suppression of *SpPI* expression in dioecious spinach (*Spinacia oleracea*) also influences stamen initiation, often leading to the complete conversion of male flowers into female flowers (Pfent *et al*., 2005; Sather *et al*., 2010). Thus, it seems likely that *QoPI* may be involved in the initiation of stamen primordia, and the disruption of the typical expression dynamics of this gene during female flower development may underlie the formation of hermaphroditic flowers.

In *Q. suber* and *C. sativa*, the AP3 and TM6 proteins were found to interact with the respective PI homologues, suggesting that the canonical obligate interaction of AP3- and PI-like proteins, necessary for the fulfilment of the B-function in stamen development, is conserved (Riechmann *et al*., 1996; Lenser *et al*., 2009; Sobral and Costa, 2017; Alhinho *et al*., 2021). Considering the high degree of sequence conservation within the Fagaceae, it is likely that these protein-protein interactions have been retained in *Q. robur* and *Q. orocantabrica*. As such, the restriction of *QoPI* expression to flowers that present stamens suggests that this obligate heterodimerization may also be conserved in these species.

The C-class gene *QoAG* is expressed in both male and female flowers, in similarity with other Fagaceae (Sobral and Costa, 2017; Alhinho *et al*., 2021; Liu *et al*., 2021). By contrast, *QoSHP* was found to be exclusive to female and hermaphroditic flowers, a trait that differs from *Q. suber*, *C. sativa* and *C. mollissima*. In these species, the *SHP* homologue is expressed in both flowers and is thought to fulfil the C-class role instead of the canonical C-class *AG* (Sobral and Costa, 2017; Alhinho *et al*., 2021; Liu *et al*., 2021). However, the apparent absence of *QoSHP* expression in *Q. orocantabrica* male flowers may indicate that *QoAG* is sufficient to fulfil the C-class function in the stamen whorl, and *QoSHP* may function as a D-class gene. These results highlight a potential functional divergence and plasticity of C/D-class genes within the Fagaceae family, as was reported for the wider eudicots following the C/D-family gene duplications (Causier *et al*., 2005; Fourquin and Ferrándiz, 2012).

In general, the results here presented indicate that the dynamics of floral sex identity in oaks is more plastic than traditionally assumed. The association between recurrent hermaphroditism, secondary flowering and polycyclic growth suggests that oak unisexuality may be a dominant, but not exclusive developmental outcome, dependant on the tight regulation of floral homeotic genes. Future studies across additional white-oak lineages and biogeographic regions will be essential to determine whether this latent floral plasticity is widespread within the genus, or whether it is more frequent in environmentally heterogeneous refugial systems such as the Iberian Peninsula, where ecological transitions, recurrent summer drought, thermal extremes and extended growing seasons may favour phenological instability and reproductive flexibility.

## Author Contributions

H.R.A. and M.M. were involved in sample collection and phenological and histological analysis. H.R.A. performed all laboratory work and with M.M.R.C. conceptualized and designed the study. H.R.A. wrote the manuscript, with contributions from all the authors. M.M.R.C. was responsible for funding acquisition. M.M.R.C., H.A. and C.V.-V. were responsible for project administration and supervision. All the authors have read and approved the manuscript for publication.

## References

Alhinho AT, Ramos MJN, Alves S, et al., Costa MMR. 2021. The Dynamics of Flower Development in *Castanea sativa* Mill. Plants 10:1538. DOI: 10.3390/plants10081538.

Angenent GC, Colombo L. 1996. Molecular control of ovule development. Trends in Plant Science 1: 228–232. DOI: 10.1016/1360-1385(96)86900-7.

Barrett SCH. 2002. The evolution of plant sexual diversity. Nature Reviews Genetics 3: 274–284. DOI: 10.1038/nrg776.

Benlloch R, d’Erfurth I, Ferrandiz C, et al., Ratet P. 2006. Isolation of *mtpim* Proves *Tnt1* a Useful Reverse Genetics Tool in *Medicago truncatula* and Uncovers New Aspects of *AP1*-Like Functions in Legumes. Plant Physiology 142: 972–983. DOI: 10.1104/pp.106.083543.

Bobinac M, Batos B, Miljković D, Radulović S. 2012. Polycylism and phenological variability in the common oak (*Quercus robur* L.). Archives of Biological Sciences 64: 97–105.

Bobinac M, Tucović A, Isajev V. 2000. Summer flowering properties of pedunculate oak and Virgilius’s oak. Bulletin of the Faculty of Forestry 83: 55–65.

Borgardt SJ. 2013. Pistillate floral and fruit development in *Quercus* subgenus *Quercus* (Fagaceae). PhD Thesis, Faculty of the Graduate School of Cornell University.

Borzan Ž. 2000. Hermaphroditic flowering in the “Green oak”, growing in Northern Dalmatia, Croatia. Glasnik za Šumske Pokuse 37: 425–439.

Borzan Ž, Stabentheiner E. 2002. Biological and taxonomical investigations of some oak species. Acta Botanica Croatica 61: 135–144.

Bowman JL, Moyroud E. 2024. Reflections on the ABC model of flower development. The Plant Cell 36: 1334–1357. DOI: 10.1093/plcell/koae044.

Bowman JL, Smyth DR, Meyerowitz EM. 1989. Genes directing flower development in Arabidopsis. The Plant Cell 1: 37–52. DOI: 10.1105/tpc.1.1.37.

Bowman JL, Smyth DR, Meyerowitz EM. 1991. Genetic interactions among floral homeotic genes of *Arabidopsis*. Development 112: 1–20. DOI: 10.1242/dev.112.1.1.

Burger WC. 1975. The species concept in *Quercus*. Taxon 24: 45–50. DOI: 10.2307/1218998.

Camus A. 1936–1938. *Les Chénes: Monographie du genre Quercus*: *Tome I. Genre Quercus, sous-genre Cyclobalanopsis, sous-genre Euquercus (sections Cerris et Mesobalanus)*. Paris: Paul Lechevalier.

Causier B, Castillo R, Zhou J, et al., Davies B. 2005. Evolution in action: following function in duplicated floral homeotic genes. Current Biology 15: 1508–1512. DOI: 10.1016/j.cub.2005.07.063.

Chang S, Puryear J, Cairney J. 1993. A simple and efficient method for isolating RNA from pine trees. Plant Molecular Biology Reporter 11: 113–116. DOI: 10.1007/BF02670468.

Charlesworth B, Charlesworth D. 1978. A model for the evolution of dioecy and gynodioecy. The American Naturalist 112: 975–997. DOI: 10.1086/283342.

Coen ES, Meyerowitz EM. 1991. The war of the whorls: genetic interactions controlling flower development. Nature 353: 31–37. DOI: 10.1038/353031a0.

Colombo L, Franken J, Koetje E, et al., van Tunen AJ. 1995. The petunia MADS box gene *FBP11* determines ovule identity. The Plant Cell 7: 1859–1868. DOI: 10.1105/tpc.7.11.1859.

Coombes A. 2025, January 14. Oaks behaving strangely. Allen Coombes’s Blog, International Oak Society. https://www.internationaloaksociety.org/content/oaks-behaving-strangely.

Denk T, Grimm GW, Manos PS, Deng M, Hipp AL. 2017. An updated infrageneric classification of the oaks: Review of previous taxonomic schemes and synthesis of evolutionary patterns. In: Gil-Pelegrín E, Peguero-Pina J, Sancho-Knapik D, eds. Oaks: Physiological ecology. Exploring the functional diversity of genus Quercus L. Cham: Springer, 13–38. DOI: 10.1007/978-3-319-69099-5_2.

Ditta G, Pinyopich A, Robles P, Pelaz S, Yanofsky MF. 2004. The *SEP4* gene of *Arabidopsis thaliana* functions in floral organ and meristem identity. Current Biology 14: 1935–1940. DOI: 10.1016/j.cub.2004.10.028.

Ducousso A, Michaud H, Lumaret R. 1993. Reproduction and gene flow in the genus *Quercus* L. Annales des Sciences Forestières 50: 91–106. DOI: 10.1051/forest:19930708.

Edgar RC. 2004. MUSCLE: multiple sequence alignment with high accuracy and high throughput. Nucleic Acids Research 32: 1792–1797. DOI: 10.1093/nar/gkh340.

Elena-Rossello JA, de Rio JM, Valdecantos JLG, Santamaria IG. 1993. Ecological aspects of the floral phenology of the cork-oak (*Q. suber* L): why do annual and biennial biotypes appear? Annals of Forest Science 50: 114–121. DOI: 10.1051/forest:19930710.

Fourquin C, Ferrándiz C. 2012. Functional analyses of AGAMOUS family members in *Nicotiana benthamiana* clarify the evolution of early and late roles of C-function genes in eudicots. The Plant Journal 71: 990–1001. DOI: 10.1111/j.1365-313X.2012.05046.x.

Franco JA. 1990. *Quercus*. In: Castroviejo S, Laínz M, González GL, et al., Villar L, eds. Flora Iberica. Vol. 2. Madrid: Real Jardín Botánico, CSIC, 15–36.

García-Mozo H, López-Orozco R, Oteros J, Galán C. 2022. Factors driving autumn *Quercus* flowering in a thermos-Mediterranean area. Agronomy 12: 2596. DOI: 10.3390/agronomy12112596.

Greene EL. 1889. *Illustrations of west American oaks*. San Francisco: Bosqui Engraving and Printing Company.

Hipp AL, Manos PS, Hahn M, et al., Valencia-Avalos S. 2020. Genomic landscape of the global oak phylogeny. New Phytologist 226: 1198–1212. DOI: 10.1111/nph.16162.

Irish VF, Sussex IM. 1990. Function of the *apetala-1* gene during *Arabidopsis* floral development. The Plant Cell 2: 741–753. DOI: 10.1105/tpc.2.8.741.

Jackson DP. 1991. In situ hybridization in plants. In: Bowles DJ, Gurr SJ, McPhereson M, eds. Molecular Plant Pathology: A Practical Approach. Vol. 1. Oxford: Oxford University Press, 163–174.

Kaul RB. 1985. Reproductive morphology of *Quercus* (Fagaceae). American Journal of Botany 72: 1962–1977. DOI: 10.1002/j.1537-2197.1985.tb08470.x.

Kaul RB. 1986. Evolution and reproductive biology of inflorescences in *Lithocarpus*, *Castanopsis*, *Castanea* and *Quercus* (Fagaceae). Annals of the Missouri Botanical Garden 73: 284–296. DOI: 10.2307/2399114.

Keuter A. 2018. Observations of hermaphroditic late-season flowering in the red oak *Quercus agrifolia*. Phytoneuron 2018: 19.

Kramer EM, Dorit RL, Irish VF. 1998. Molecular evolution of genes controlling petal and stamen development: duplication and divergence within the *APETALA3* and *PISTILLATA* MADS-box gene lineages. Genetics 149: 765–783. DOI: 10.1093/genetics/149.2.765.

Kramer EM, Irish VF. 2000. Evolution of the petal and stamen developmental programs: evidence from comparative studies of the lower eudicots and basal angiosperms. International Journal of Plant Sciences 161: S29–S40. DOI: 10.1086/317576.

Kramer E, Jaramillo MA, Di Stilio VS. 2004. Patterns of gene duplication and functional evolution during the diversification of the *AGAMOUS* subfamily of MADS box genes in angiosperms. Genetics 166: 1011–1023. DOI: 10.1093/genetics/166.2.1011.

Kubitzki K. 1993. Fagaceae. In: Kubitzki K, Rohwer JG, Bittrich V, eds. The Families and Genera of Vascular Plants. Vol. 2. Berlin: Springer, 301–309.

Le Provost G, Herrera R, Paiva JA, Chaumeil P, Salin F, Plomion C. 2007. A micromethod for high throughput RNA extraction in forest trees. Biological Research 40: 291–297. DOI: 10.4067/S0716-97602007000400003.

Lenser T, Theißen G, Dittrich P. 2009. Developmental robustness by obligate interaction of class B floral homeotic genes and proteins. PLOS Computational Biology 5: e1000264. DOI: 10.1371/journal.pcbi.1000264.

Liu Y, Chen G, Gao Y, et al., Su S. 2021. Identification and characterization of MADS-box genes involved in floral organ development in Chinese chestnut (*Castanea mollissima* Blume). Horticultural Science and Technology 39: 482–496. DOI: 10.7235/HORT.20210043.

Livak KJ, Schmittgen TD. 2001. Analysis of relative gene expression data using real-time quantitative PCR and the 2^-ΔΔCT^ method. Methods 25: 402–408. DOI: 10.1006/meth.2001.1262.

Maqueda SR. 2007. Anomalías en las estructuras reproductoras del género *Quercus* L. (Fagaceae). Folia Botanica Extremadurensis 1: 75–78.

Marum L, Miguel A, Ricardo CP, Miguel C. 2012. Reference gene selection for quantitative real-time PCR normalization in *Quercus suber*. PLOS One 7: e35113. DOI: 10.1371/journal.pone.0035113.

Martínez C, Manzano S, Megías Z, et al., Jamilena M. 2014. Molecular and functional characterization of *CpACS27A* gene reveals its involvement in monoecy instability and other associated traits in squash (*Cucurbita pepo* L.). Planta 239: 1201–1215. DOI: 10.1007/s00425-014-2043-0.

de Martino G, Pan I, Emmanuel E, Levy A, Irish VF. 2006. Functional analyses of two tomato *APETALA3* genes demonstrate diversification in their roles in regulating floral development. The Plant Cell 18: 1833–1845. DOI: 10.1105/tpc.106.042978.

Merkle SA, Feret PP, Croxdale JG, Sharik TL. 1980. Development of floral primordia in white oak. Forest Science 26: 238–250. DOI: 10.1093/forestscience/26.2.238.

Meyerowitz EM, Bowman JL, Brockman LL, et al., Weigel D. 1991. A genetic and molecular model for flower development in *Arabidopsis thaliana*. Development 113: 157–167. DOI: 10.1242/dev.113.Supplement_1.157.

Mitchell CH, Diggle PK. 2005. The evolution of unisexual flowers: morphological and functional convergence results from diverse developmental transitions. American Journal of Botany 92: 1068–1076. DOI: 10.3732/ajb.92.7.1068.

Natividade J. 1950. Subericultura. Lisboa: Ministerio da Economia, Direçao Geral dos Servicios Florestais e Aquicolas.

Nixon KC. 1993. Infrageneric classification of *Quercus* (Fagaceae) and typification of sectional names. Annals of Forest Science 50: 25s–34s. DOI: 10.1051/forest:19930701.

Pelaz S, Ditta GS, Baumann E, Wisman E, Yanofsky MF. 2000. B and C floral organ identity functions require *SEPALLATA* MADS-box genes. Nature 405: 200–203. DOI: 10.1038/35012103.

Plomion C, Aury JM, Amselem J, et al., Kremer A. 2016. Decoding the oak genome: public release of sequence data, assembly, annotation and publication strategies. Molecular Ecology Resources 16: 254–265. DOI: 10.1111/1755-0998.12425.

Pfent C, Pobursky KJ, Sather DN, Golenberg EM. 2005. Characterization of *SpAPETALA3* and *SpPISTILLATA*, B class floral identity genes in *Spinacia oleracea*, and their relationship to sexual dimorphism. Development Genes and Evolution 215: 132–142. DOI: 10.1007/s00427-004-0459-4.

Rajatiya JH, Varu DK, Halepotara FH, Solanki MB. 2018. Correlation of climatic parameters with flowering characters of mango. International Journal of Pure and Applied Bioscience 6: 597–601. DOI: 10.18782/2320-7051.6523.

Rangel SR, Zenteno ECR, Maqueda SG. 2000. Flores hermafroditas de *Quercus glaucoides* Mart. & Gal. (Fagaceae) en el Estado de Michoacán, México. Acta Botanica Mexicana 52: 49–54. DOI: 10.21829/abm52.2000.855.

Riechmann JL, Krizek BA, Meyerowitz EM. 1996. Dimerization specificity of *Arabidopsis* MADS domain homeotic proteins APETALA1, APETALA3, PISTILLATA, and AGAMOUS. Proceedings of the National Academy of Sciences of the United States of America 93: 4793–4798. DOI: 10.1073/pnas.93.10.4793.

Romero S, Rojas EC, Garay-Velázquez OH. 2007. Presencia de flores hermafroditas en *Quercus rugosa* (Fagaceae) en el Estado de México (México). Anales del Jardín Botánico de Madrid 64: 223–227. DOI: 10.3989/ajbm.2007.v64.i2.179.

Sather DN, Jovanovic M, Golenberg EM. 2010. Functional analysis of B and C class floral organ genes in spinach demonstrates their role in sexual dimorphism. BMC Plant Biology 10: 46–54. DOI: 10.1186/1471-2229-10-46.

Serrazina S, Santos C, Machado H, et al., Costa R. 2015. *Castanea* root transcriptome in response to *Phytophthora cinnamomi* challenge. Tree Genetics & Genomes 11: 6. DOI: 10.1007/s11295-014-0829-7.

Schwarz O. 1937. Monographie der Eichen Europas und des Mittelmeergebietes. Repertorium novarum speciarum regni vegetabilis 1: 1–200.

Sobral R, Costa MMR. 2017. Role of floral organ identity genes in the development of unisexual flowers of *Quercus suber* L. Scientific Reports 7: 10368. DOI: 10.1038/s41598-017-10732-0.

Sobral R, Silva HG, Laranjeira S, Magalhães J, Andrade L, Alhinho AT, Costa MMR. 2020. Unisexual flower initiation in the monoecious *Quercus suber* L.: a molecular approach. Tree Physiology 40: 1260–1276. DOI: 10.1093/treephys/tpaa061.

Soepadmo E, van Steenis CGGJ. 1972. Fagaceae. In: Flora Malesiana Series 1, Spermatophyta. 7: 265–403.

Tamura K, Stecher G, Sudhir K. 2021. MEGA11: molecular evolutionary genetics analysis version 11. Molecular Biology and Evolution 38: 3022–3027. DOI: 10.1093/molbev/msab120.

Theißen G. 2001. Development of floral organ identity: stories from MADS house. Current Opinion in Plant Biology 4: 75–85. DOI: 10.1016/S1369-5266(00)00139-4.

Theißen G, Melzer R, Rümpler F. 2016. MADS-domain transcription factors and the floral quartet model of flower development: linking plant development and evolution. Development 143: 3259–3271. DOI: 10.1242/dev.134080.

Tucker JM. 1972. Hermaphroditic flowers in Californian oaks. Madroño 21: 482–486.

Tucker JM, Neilson RP, Wullstein LH. 1980. Hermaphroditic flowering in Gambel oak. American Journal of Botany 67: 1265–1267. DOI: 10.1002/j.1537-2197.1980.tb07759.x.

Varela MC, Valdiviesso T. 1996. Phenological phases of *Quercus suber* L. flowering. Forest Genetics 3: 93–102.

Vila-Viçosa C, Arenas-Castro S, Marcos B, et al., Gonçalves J. 2020. Combining satellite remote sensing and climate data in species distribution models to improve the conservation of Iberian white oaks (*Quercus* L.). ISPRS International Journal of Geo-Information 9: 735. DOI: 10.3390/ijgi9120735.

Vila-Viçosa C, Arraiano-Castilho R, Vázquez FM, et al., Azevedo H. 2026. The Iberian white-oak syngameon as a legacy of introgression in southern Europe. bioRxiv 730892. DOI: 10.64898/2026.06.09.730892.

Vila-Viçosa C, Capelo J, Alves P, Almeida R, Vázquez FM. 2023. New annotated checklist of Portuguese oaks (*Quercus*, Fagaceae). Mediterranean Botany 44: e79286. DOI: 10.5209/mbot.79286.

Vrebalov J, Ruezinsky D, Padmanabhan V, et al., Giovannoni J. 2002. A MADS-box gene necessary for fruit ripening at the tomato *Ripening-Inhibitor* (*Rin*) locus. Science 296: 343–346. DOI: 10.1126/science.1068181.

Zhang C, Sun Y, Yu X, Li H, Bao M, He Y. 2021. Functional conservation and divergence of five AP1/FUL-like genes in marigold (*Tagetes erecta* L.). Genes 12: 2011. DOI: 10.3390/genes12122011.

