## Supplementary Material for "Recurrent Hermaphroditism and Sex-Biased ABCDE Gene Expression Reveal Latent Floral Plasticity in the Pedunculate Oak Lineage (*Q. robur* s.l.)"

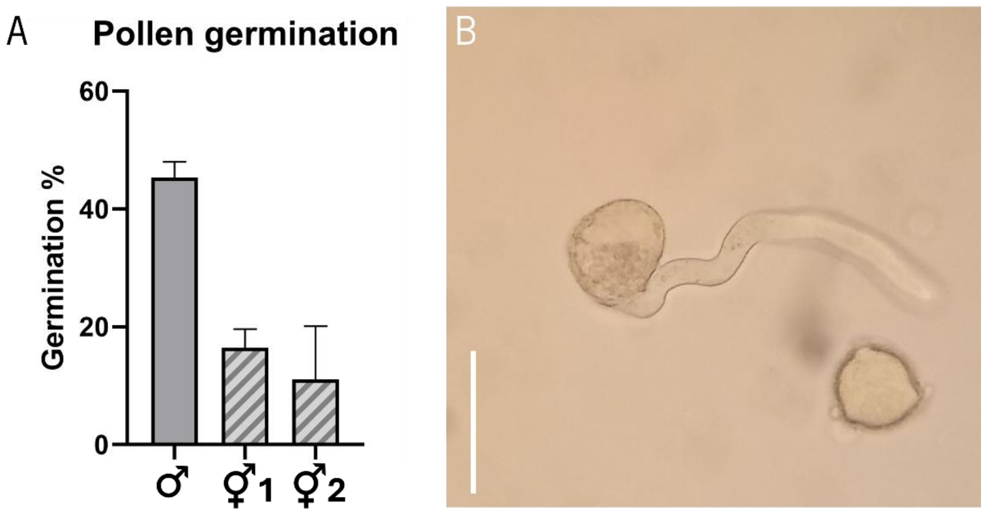


**Fig. S1: Pollen germination assay. A)** Pollen germination rate from male flowers (♂), and hermaphroditic flowers from a first (⚥1) and second (⚥2) flowering events. Germination rate was calculated from three technical replicates, with a minimum of 500 pollen grains counted in each replicate; **B)** Germinating pollen grain from a hermaphroditic flower. Scale bar: 50 µm.

**A**

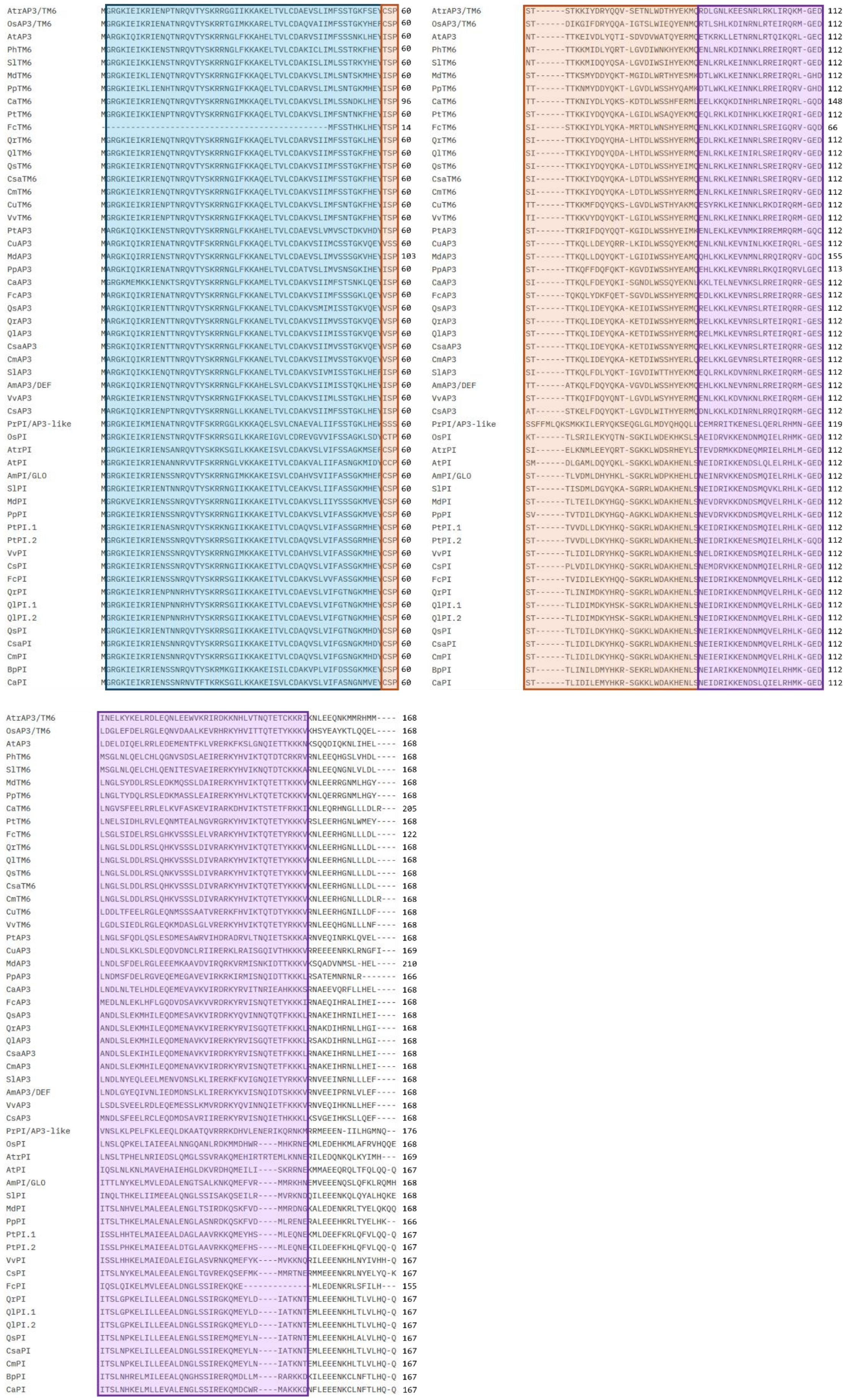

**B**

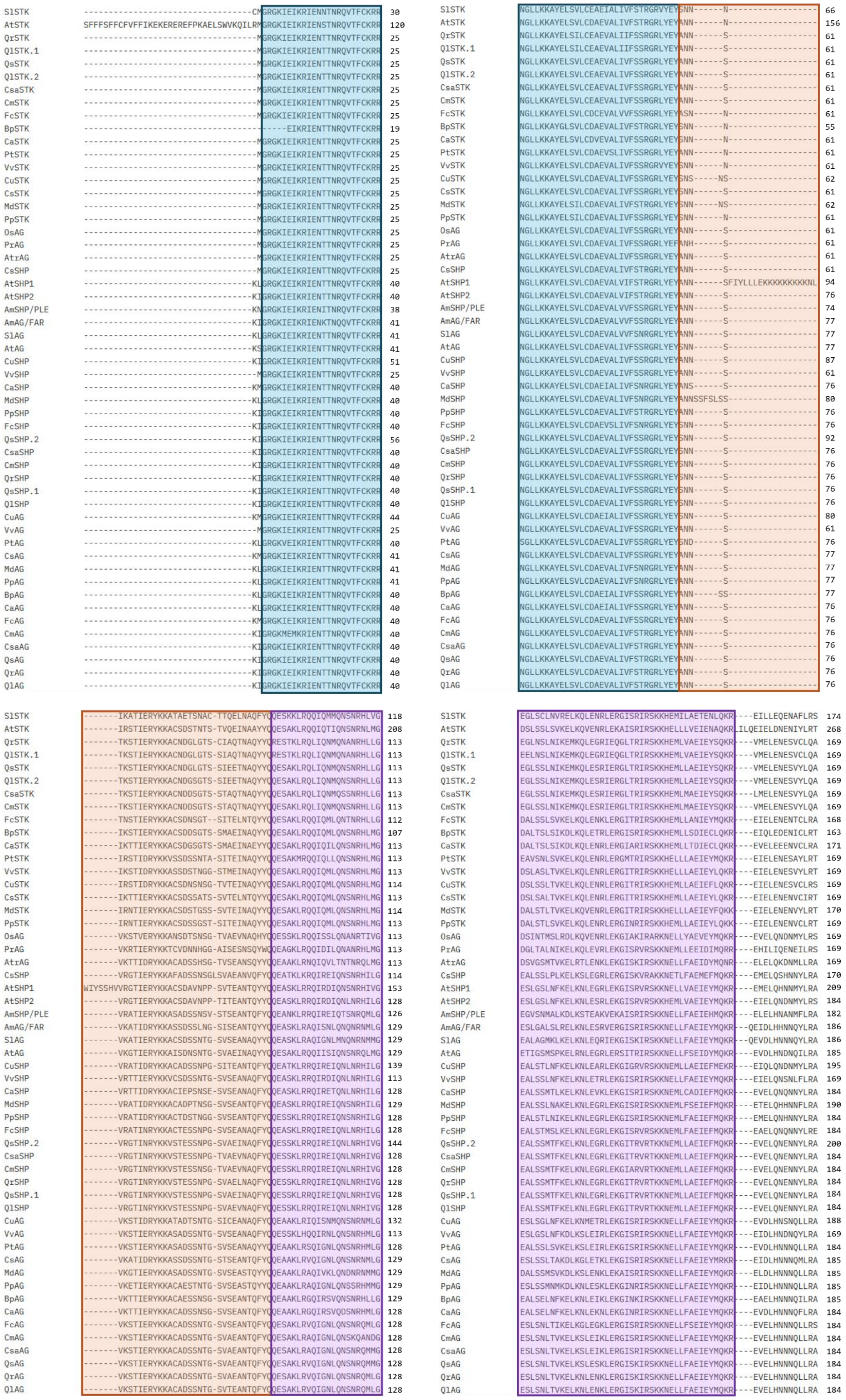

**Fig. S2: Protein alignment of BCD-like sequences. A)** B-class proteins; **B)** C and D-class proteins. Blue boxes: MADS-domain; Orange boxes: I-domain; Purple boxes: K-domain.

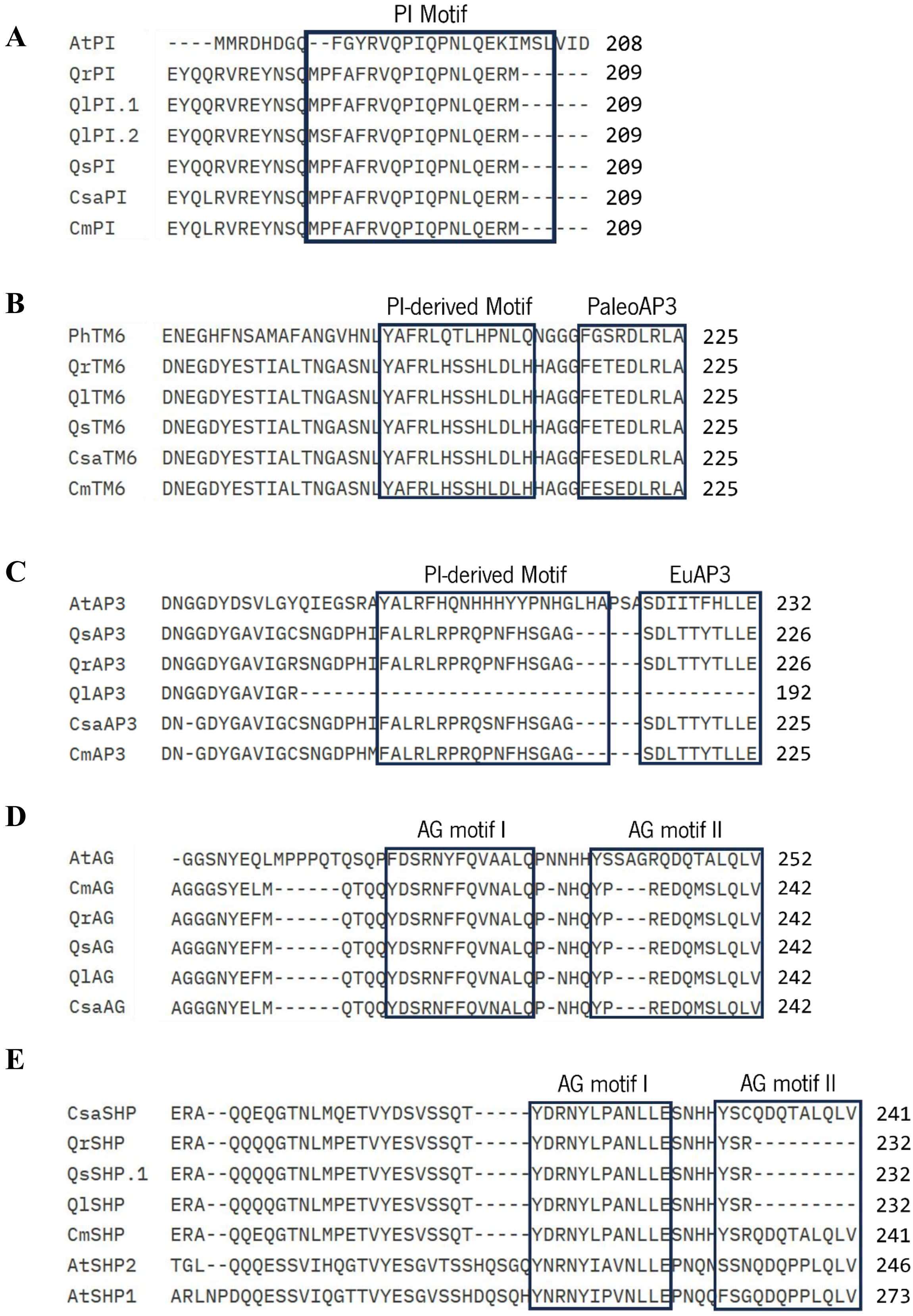


**Figure S3: Motif comparison of the C-terminal domains of Fagaceae BCD-like proteins. A)** PI-like proteins; **B)** TM6-like proteins; **C)** AP3-like proteins; **D)** AG-like proteins; **E)** SHP-like proteins.

**Table S1** – List of primers used for quantitative RT-PCR

| Amplicon | Direction | Sequence | Annealing Temperature (°C) | Fragment size (bp) |
|---|---|---|---|---|
| *QoPP2AA3* | Forward | GCAGCACATAATTCCACAGGTT | 60 | 236 |
| | Reverse | TCTCCACCACAGACTGATCAAC | | |
| *QoEF1a* | Forward | CATCATGAACCACCCGGTCA | 60 | 168 |
| | Reverse | CCCAGCATCTCCGTTCTTCA | | |
| *QoAP3* | Forward | GACCGCAAGTACCAGGTGAT | 60 | 213 |
| | Reverse | CCTAGGTCTCAGACGCAAAG | | |
| *QoPI* | Forward | CCGAGAAATGCAGATGGAGT | 60 | 199 |
| | Reverse | AATAGGCTGCACACGGAAGG | | |
| *QoTM6* | Forward | CTGCTCGATCTTAGGGCAAG | 60 | 103 |
| | Reverse | AGTTGGAGGCACCATTTGTC | | |
| *QoAG* | Forward | CCAGCTTCTCCGAGCAAAGA | 60 | 175 |
| | Reverse | CATCTGGTCTTCACGTGGGT | | |
| *QoSHP* | Forward | AGGGAAGTTGAGCGCAAAA | 60 | 151 |
| | Reverse | CTGGGAGGTAGTTCCGATCA | | |

**Table S2** – List of protein accession numbers

| Protein | Accession Number | | Protein | Accession Number |
|---|---|---|---|---|
| **B-class Family** | | | | |
| QrPI | XP_050255112 | | CaTM6 | XP_059439080 |
| QsPI | XP_023887186 | | MdTM6 | KAM0970310 |
| QlPI1 | XP_030933539 | | PpTM6 | XP_007202502 |
| QlPI2 | XP_030933547 | | PtTM6 | XP_024456101 |
| CsaPI | XP_075650620 | | CuTM6 | GAY57755 |
| CmPI | KAF3974022 | | PhTM6 | AAF73933 |
| FcPI | GMY06659 | | VvTM6 | NP_001267937 |
| BpPI | CAD32764 | | SlTM6 | NP_001311309 |
| CaPI | XP_059454186 | | QrAP3 | XP_050267596 |
| CsPI | NP_001292651 | | QsAP3 | XP_023924416 |
| MdPI | CAC28022 | | QlAP3 | XP_030933764 |
| PpPI | XP_020410381 | | CsaAP3 | XP_075652509 |
| PtPI1 | XP_002307460 | | CmAP3 | KAF3962544 |
| PtPI2 | XP_002300964 | | FcAP3 | GMY28324 |
| AtPI | AT5G20240 | | CaAP3 | XP_059461662 |
| VvPI | NP_001267875 | | CsAP3 | NP_001295864 |
| SlPI | NP_001234075 | | MdAP3 | XP_028962429 |
| AmPI | Q03378 | | PpAP3 | XP_020415740 |
| OsPI | AAC05723 | | PtAP3 | XP_006386194 |
| AtrPI | XP_006847167 | | CuAP3 | GAY42987 |
| PrAP3/PI | AAF28863 | | AtAP3 | AT3G54340 |
| QrTM6 | XP_050292299 | | VvAP3 | XP_002279735 |
| QsTM6 | XP_023911257 | | SlAP3 | NP_001234077 |
| QlTM6 | XP_030975573 | | AmAP3 | BAI68389 |
| CsaTM6 | XP_075656191 | | OsAP3 | BAH22555 |
| CmTM6 | KAF3962544 | | AtrAP3 | XP_011628954 |
| FcTM6 | GMY12162 | | | |
| **C-class Family** | | | | |
| QrAG | XP_050290781 | | AtrAG | NP_001292764 |
| QsAG | XP_023894765 | | PrAG | AAD09342 |
| QlAG | XP_030971533 | | QrSHP | XP_050249327 |
| CsaAG | XP_075667291 | | QsSHP.1 | XP_023917960 |
| CmAG | AAZ77747 | | QsSHP.2 | POF03476 |
| FcAG | GMY37787 | | QlSHP | XP_030932117 |
| BpAG | CAB95649 | | CsaSHP | XP_075671509 |
| CaAG | XP_059438412 | | CmSHP | UYO08119 |
| CsAG | NP_001292633 | | FcSHP | GMY08913 |
| MdAG | XP_008383546 | | CaSHP | XP_059450236 |
| PpAG | XP_007211925 | | CsSHP | NP_001292697 |
| PtAG | XP_024455023 | | MdSHP | XP_070672366 |
| CuAG | BAF34911 | | PpSHP | XP_007217264 |
| AtAG | AT4G18960 | | CuSHP | BAF34914 |
| VvAG | NP_001268097 | | AtSHP.1 | AT3G58780 |
| SlAG | NP_001295225 | | CsSHP | NP_001292697 |
| AmAG | BAI68392 | | MdSHP | XP_070672366 |
| OsAG | ACY26070 | | PpSHP | XP_007217264 |

| Protein | Accession Number | | Protein | Accession Number |
|---|---|---|---|---|
| CuSHP | BAF34914 | | BpSTK | CAK55150 |
| AtSHP.1 | AT3G58780 | | CaSTK | XP_059449028 |
| AtSHP.2 | AT2G42830 | | CsSTK | NP_001267506 |
| VvSHP | NP_001268105 | | MdSTK | NP_001280931 |
| AmSHP | AAB25101 | | PpSTK | XP_020416253 |
| QrSTK | XP_050240882 | | PtSTK | XP_006376118 |
| QsSTK | XP_065624585 | | CuSTK | GAY61853 |
| QlSTK | XP_030975845 | | AtSTK | AT4G09960 |
| CsaSTK | XP_075655610 | | VvSTK | QSX80212 |
| CmSTK | KAF3963586 | | SlSTK | XP_004241906 |
| FcSTK | GMY09374 | | | |
